# Distinct generation of subjective vividness and confidence during naturalistic memory retrieval in angular gyrus

**DOI:** 10.1101/2021.03.10.434526

**Authors:** Futing Zou, Sze Chai Kwok

## Abstract

Our subjective experience of remembering guides and monitors the reconstruction of past and simulation of the future, which enables us to identify mistakes and adjust our behavior accordingly. However, it remains incompletely understood what underlies the process of subjective mnemonic experience. Here, we combined behavior, repetitive transcranial magnetic stimulation (rTMS), and functional neuroimaging to probe whether vividness and confidence are generated differently during retrieval. We found that pre-retrieval rTMS targeting the left angular gyrus (AnG) selectively attenuated the vividness efficiency compared to control stimulation while keeping metacognitive efficiency and objective memory accuracy unaffected. Using trial-wise data, we showed that AnG stimulation altered the mediating role of vividness in confidence in the accuracy of memory judgment. Moreover, resting-state functional connectivity of hippocampus and AnG was specifically associated with vividness efficiency, but not metacognitive efficiency across individuals. Together, these results identify the causal involvement of AnG in gauging the vividness, but not the confidence, of memory, thereby suggesting a differentiation account of conscious assessment of memory by functionally and anatomically dissociating the monitoring of vividness from confidence.

## Introduction

According to Endel Tulving (Tulving, 1972, 1985), the conception of episodic memory is identified with autonoetic awareness, which gives rise to remembering of personally experienced events. The process of explicitly remembering a specific previous event is often accompanied by a subjective sense of recollection, which enables us to monitor experiences, identify mistakes, and guide future behavior accordingly. It is therefore crucial to understand what underlies the subjective mnemonic experiences, such as subjective vividness of the memory and confidence in the memory decisions. In memory research, vividness and confidence are often used interchangeably under the umbrella of “subjective experience”. However, an important and intriguing idea is that the processes of generating vividness and confidence operate differently during memory retrieval. Specifically, confidence is often used as a measure of the capacity to evaluate one’s own cognitive processes, referred to as metacognition (Metcalfe, 1997), which has been studied across a range of task domains, including perceptual, memory, social, and value-based decisions (Bang et al., 2017; De Martino et al., 2012; McCurdy et al., 2013; Morales et al., 2018; Ye et al., 2018). By comparison, vividness is a relatively specific measure of episodic memory recollection, which has been used to assess the degree to which the retrieved content is rich and detailed. On this basis, we reasoned that the computation of vividness should be partially, if not fully, independent from confidence of memory.

In this way, vividness and confidence of memory could be mediated by distinct neural mechanisms and even in different brain regions. A candidate region thought to differently support these two subjective mnemonic components is left lateral parietal cortex, in particular the angular gyrus (AnG). The left AnG is widely thought to play an important role in subjective experience of remembering. For example, a number of human neuroimaging studies have shown that activity in AnG is associated with subjective reports of vividness (Bonnici et al., 2016; Kuhl & Chun, 2014) and confidence (Qin et al., 2011) during episodic memory retrieval. Consistently, disruption of left AnG processing by transcranial magnetic stimulation (TMS) has been found to selectively reduce confidence but leaving objective retrieval success intact (Wynn et al., 2018; Yazar et al., 2014; but also see Branzi et al., 2021). These results, however, have focused primarily on the level of confidence or vividness rating during memory retrieval, thereby leaving unanswered whether this region supports the ability to faithfully monitor subjective sense of remembering (i.e., the correspondence between objective memory performance and subjective memory reports). Furthermore, the left AnG has been proposed to be involved in the integration of mnemonic features into a conscious representation that enables the subjective experience of remembering (Bonnici et al., 2016; Humphreys et al., 2021), which is analogous with the definition of the vividness of memory rather than a general subjective sense of confidence in memory decisions. While it is difficult to completely rule the AnG out in confidence processing, we are inclined to theories asserting that confidence signal is modulated by meta-level information, above and beyond integration of multisensory information (De Martino et al., 2012; Shekhar & Rahnev, 2018). To our knowledge, no study has provided evidence for the involvement of AnG in metacognitive processing. We thus reasoned that the left AG might exhibit disproportional engagement in the computation of vividness relative to confidence. It is important to note that we are interested in the degree to which confidence and vividness are related to objective memory performance, namely, metacognitive (confidence) efficiency and vividness efficiency, instead of the level of subjective ratings.

Here we aimed to ask two key questions: i) Are vividness and confidence dissociable subjective components during episodic memory? ii) Does the AnG support the subjective assessment of memory quality? We addressed these questions by using both TMS and MRI methods. Specifically, to temporarily manipulate AnG function, we administered an inhibitory repetitive TMS protocol to left AnG as well as to a control site (vertex) in a within-subjects design. Following a 20-min rTMS protocol, we asked participants to report the vividness of mental replay of a target scene before the memory judgments and confidence ratings. Of note, to ensure that vividness and confidence ratings are based on the same segment of a piece of memory within a trial, we set to test participants’ objective memory that largely depends upon the quality of the preceding mental replay. Accordingly, in the memory judgments, participants were asked to perform a temporal proximity judgment between two scenes with respect to the target scene. The temporal proximity judgment task requires participants to compare the temporal distance of two chunks of a specific episode, which demands participants to mentally replay the cue related scenes for successful memory retrieval. Given the nature of our temporal proximity task, a correct memory response will be dependent on precise recollection of all three of these scenes. We expected that recollection is in turn related to participant’s subjective evaluation of recall (i.e., subjective vividness). For an accurate comparison between these subjective experiences, we quantified the efficiencies of the two subjective memory ratings by computing the trial-by-trial correspondence between objective memory performance and subjective reports (Fleming & Daw, 2016; Maniscalco & Lau, 2012). As the correspondence between objective and subjective memory reports increases, subjective awareness of memory approaches ideal. Given the known involvement of hippocampus in memory recollection and the richness of re-experiencing (Ford & Kensinger, 2016; Gilboa et al., 2004), we also employed a functional connectivity approach to assess the relationship between functional architecture of these regions and both subjective evaluation abilities. If the aforementioned hypothesis is true, we would expect to see a dissociation between vividness and metacognitive efficiency, where TMS to the left AnG will selectively affect the vividness efficiency but not metacognitive efficiency.

## Methods

### Participants

Twenty healthy young adults took part in this study (11 females and 9 males, mean age = 22.70 years, SD = 2.8, range = 18-26). The sample size was determined based on a power analysis (alpha=0.05, two-tailed, power=0.8) performed on data from our previous TMS study probing the causal role of parietal cortex on memory metacognition (Ye et al., 2018). All participants were right-handed with normal or corrected-to-normal vision, and had no contraindications for MRI or TMS. Each of them participated in two experimental sessions, giving us a within-subjects comparison to assess the influence of TMS to AnG on memory. Data from three additional participants were excluded from data analyses: one participant did not complete the experiment due to anxiety and the other two inadvertently hit the wrong response key throughout a whole test session. Participants were recruited from the East China Normal University undergraduate and graduate student population and compensated for their participation. The East China Normal University Committee on Human Research Protection approved the experimental protocol and all participants gave their written informed consent. All participants self-reported to be native Chinese speakers and had not previously seen any episodes of *Black Mirror.*

### Overview of Experimental Design

Participants completed a baseline session and two experimental sessions on separate days in a within-subjects design (Fig. 1A). Following standard MRI and TMS safety screening, participants first underwent a baseline session where structural MRI scans and resting-state fMRI scans were obtained. The structural MRI scans were used to define the subjective-specific stimulation locations and enable accurate navigation. Each experimental session consisted of two phases separated by one day: an approximately 1-hr encoding session, during which participants watched one *Black Mirror* movie, and a retrieval session one day later, during which participants received either rTMS over the left-AnG or over the vertex and completed a memory retrieval test. The retrieval began immediately after rTMS and lasted 50 min. In the retrieval phase, participants recalled relevant scenarios based on a cue image, rated their subjective vividness of the mental replay, made temporal proximity judgments, and rated their confidence of the memory judgments (Fig. 1C).

**Figure 1.**
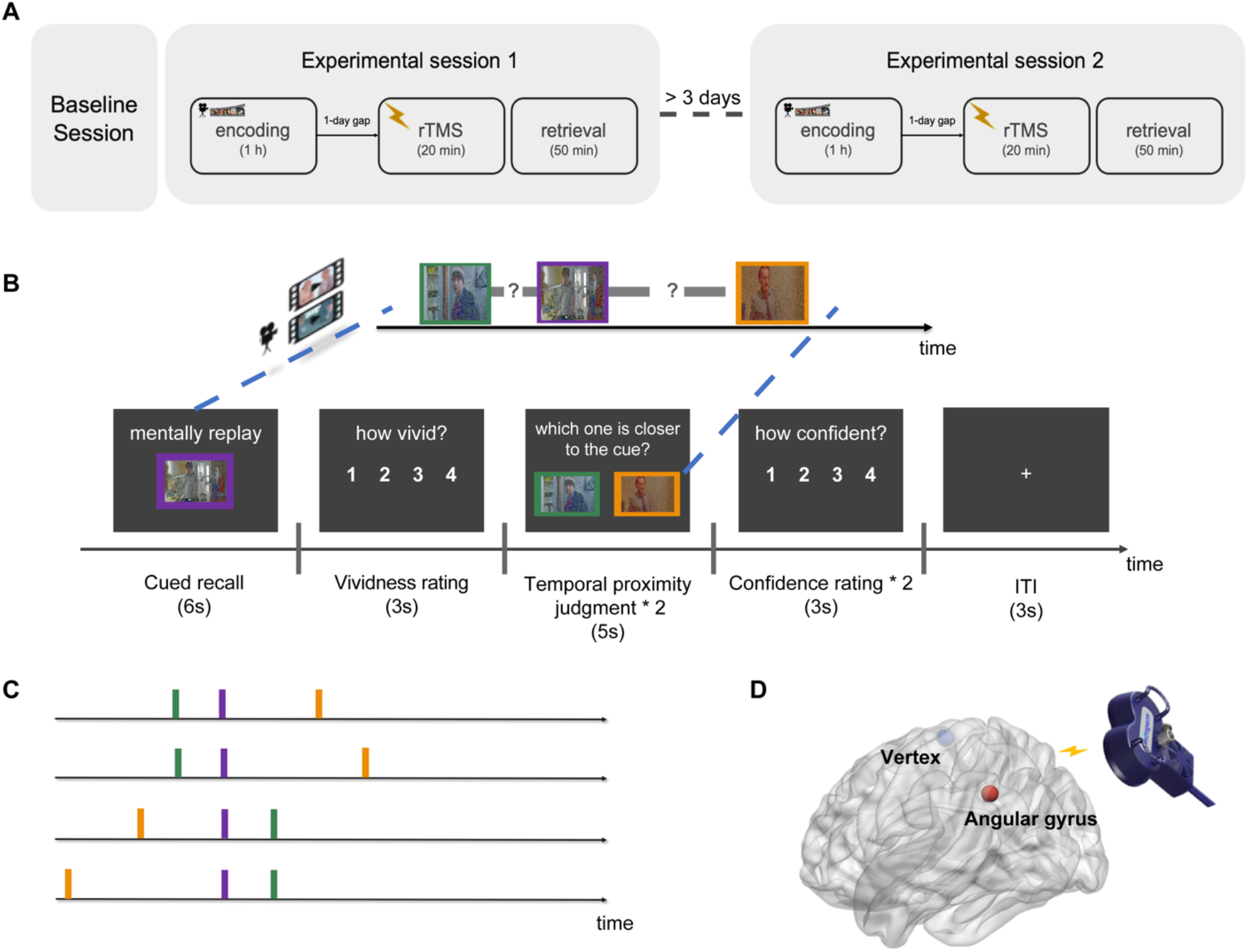
Experimental design. **(A)** Overview of task design. In each of the experimental sessions, participants viewed a 1-hr movie from *Black Mirror* at encoding. On the following day, participants received stimulation (over AnG or vertex) and performed a memory test. Movie and stimulation sites were assigned in a randomized and counterbalanced order. **(B)** Schematic overview of the memory test. Trial example: participants mentally replay related scenarios while viewing an image cue from the movie and rated the vividness of their memory. Participants were then presented with another two still frames from the movies and tested on their memory associated with the cued scene, followed by a confidence rating. Each cued recall was followed by two temporal proximity judgments. Movie stills in the figure are blurred for copyright reasons. **(C)** Triad of movie stills selection criteria (purple: cue; green: the closer one to cue; orange: the further one to cue). **(D)** Stimulation sites: AnG (red, MNI coordinate: x = −43, y = −66, z = 38) and vertex (blue, as control site).

### Memory tests

In the memory test (Fig. 1B), participants were first presented with an image cue abstracted from the movie and asked to mentally recall related scenarios in the movie as detailed as possible for 6 s. Participants were explicitly instructed to recall the full event related to the cue scene. They were instructed to replay details not only from the point of the cued scene but also those preceding the cue. This served to ensure that the corresponding vividness ratings would be related to the full event segment encompassing the cue scene. Following the mental replay, participants were allowed 3 s to rate their vividness of the memory by selecting a number from 1 to 4 (“not vivid” to “very vivid”). After the vividness rating, participants were presented with another two still frames from the movie and were asked to choose which of the two frames was temporally closer to the cue frame in the movie. On each trial, the stimulus presentation and response window lasted for 5 s. Each temporal proximity judgment was followed by a subjective confidence rating of their choice on a scale from 1 to 4 (“not confident” to “very confident”). 3 s were allowed for confidence ratings. There were two sets of temporal proximity judgment and confidence rating following each cued recall. No feedback was provided during the memory test.

### Movie scene stimuli used for encoding, cued recall, and temporal proximity judgment tests

Participants viewed two episodes of the British television series *Black Mirror* (Fig. 1B; the first episode of Season 3, *Nosedive,* and the third episode of Season 3, *Shut up and Dance)* with Chinese dubbing. Each episode was assigned to one of the experimental sessions. *Nosedive* was ~58 min long and *Shut up and Dance* was ~52 min long. For the subsequent memory retrieval test, 180 triads of still frames were extracted from each movie based on the following criteria: i) for each triad, one cue frame and two still images for temporal proximity judgments were from the adjacent scenes; ii) the absolute temporal distance between cue frame and temporally closer one to the cue was fixed. To further increase task difficulty, we selected the stimuli from four difficulty settings: hard/easy with left/right target (Fig. 1C). The occurrence of event boundaries was identified using subjective annotations. Two external observers, who did not take part in the experimental sessions of the current study and had no knowledge of the experimental design, viewed each of the movies and annotated with precision the temporal point at which they felt “a new event is starting; these are points in the movie when there is a major change in topic, location or time.” Participants were also asked to write down a short title for each event. With the participants’ boundary annotations, we looked for those boundary time points that were consistent across observers. This resulted in 50 scenes in *Nosedive* and 43 scenes in *Shut up and Dance*. Given that event boundary can affect memory retrieval (DuBrow & Davachi, 2013), this procedure allowed us to control for this potential boundary effect and equate the memory task difficulty between stimulation sites. Episode and experimental sessions were assigned in a randomized and counterbalanced order across participants.

### MRI data acquisition

Participants were scanned in a 3-tesla Siemens Trio magnetic resonance imaging scanner with a 64-channel head coil. Structural MRI images were obtained using a T1-weighted (T1w) multiecho MPRAGE protocol (field of view = 224 mm, TR = 2300 ms, TE = 2.25 ms, flip angle = 8°, voxel size = 1 × 1 × 1 mm, 192 sagittal slices) to stereotaxically guide the stimulation. Resting-state functional images were acquired with the following sequence: TR = 2450 ms, TE = 30 ms, field of view (FOV) = 192mm, flip angle = 81, voxel size = 3 × 3 × 3 mm.

### Repetitive transcranial magnetic stimulation (rTMS)

In each experimental session, participants received rTMS to either the left AnG or vertex before the memory test. The stimulation site order was counterbalanced across participants. rTMS was applied using a Magstim Rapid2 magnetic stimulator connected to a 70 mm double air film coil. The structural data obtained from each participant were used in Brainsight 2.0, a computerized frameless stereotaxic system (Rogue Research), to identify the target brain regions on a subject-by-subject basis. The stimulation sites were selected in the system by transformation of the Montreal Neurological Institute (MNI) stereotaxic coordinates to participant’s normalized brain. The sites stimulated were located in the left AnG at the MNI coordinate x=-43, y= −66, z=38, and in a control area on the vertex, which was identified at the point of the same distance to the left and the right pre-auricular, and of the same distance to the nasion and the inion (Fig. 1D). The AnG coordinate was determined from a meta-review of the parietal lobe and memory (Vilberg & Rugg, 2008). This coordinate has been adopted in several TMS studies studying subjective memory (Bonnici et al., 2016; Tibon et al., 2019; Wynn et al., 2018; Yazar et al., 2014). To target the selected stimulation sites, four fiducial points located on the face were used to co-register the anatomical MRI to the participant’s head using an infrared pointer. The real-time locations of the TMS coil and the participant’s head were monitored by an infrared camera using a Polaris Optical Tracking System (Northern Digital).

The rTMS protocol was adopted from a similar study probing episodic memory metacognition (Ye et al., 2018). This stimulation protocol has also been used to induce inhibitory effect on the AnG in a similar task (Wynn et al., 2018). Specifically, rTMS was applied at 1 Hz frequency for a continuous duration of 20 min (1200 pulses in total) at 110% of active motor threshold (MT), which was defined as the lowest TMS intensity delivered over the motor cortex necessary to elicit visible twitches of the right index finger in at least 5 of 10 consecutive pulses (Rossini et al., 2015). During stimulation, participants wore earplugs to attenuate the sound of the stimulating coil discharge. The coil was held to the scalp of the participant with a custom coil holder and the participant’s head was propped in a comfortable position. This particular stimulation magnitude and protocols of rTMS is known to induce efficacious intracortical inhibitory effects for over 60 min (Rossini et al., 2015; Thut & Pascual-Leone, 2010). Given that our task lasted 50 min, the TMS effects should have been long-lasting enough for the task. Although these inhibitory effects are known to level off within hours by the end of the stimulation, for safety reasons and to avoid carryover effects of rTMS across sessions, experimental session 1 and 2 were conducted on separate days with at least 3 days apart.

### Behavioral data analysis

Metacognition refers to one’s subjective access to their own cognitive processes, and is computed by estimating how accurate subjective ratings distinguish between correct and incorrect responses. For comparability with previous metacognition work (for review, see Fleming & Lau, 2014), we estimated memory metacognitive ability using the confidence ratings. To assess whether participants’ confidence ratings were reliably related to their objective memory performance, we computed meta-d’, a metric that quantifies the metacognitive sensitivity and is independent of confidence bias, using a Bayesian modelbased method (Fleming, 2017; Fleming & Lau, 2014). Given the metric, meta-d’, is expressed in the same units as d’, it allows a direct comparison between objective performance and metacognitive sensitivity. For example, if meta-d’ equals d’, it means that the observer is metacognitively ideal. Meta-d’ greater or less than d’ indicates metacognition that is better or worse, respectively, than the expected given task performance. Here we assessed metacognitive efficiency using the ratio meta-d’/d’, which indexes participant’s metacognitive efficiency while adjusting for the influence of objective memory performance and response bias. Similarly, to quantify the extent to which participants’ vividness ratings tracked their objective memory performance, we applied the same single-subject Bayesian meta-d’ algorithm but to vividness ratings and computed a metric termed vividness efficiency (vivid-d’/d’).

### Resting-state functional connectivity analysis

For connectivity analysis of resting-state data, resting-state functional data were first converted to Brain Imaging Data Structure (BIDS) format and verified using the BIDS validator. Data preprocessing was performed using fMRIPrep (Esteban et al., 2019) with the default processing steps, including skull stripping, motion correction, brain tissue segmentation, slice time correction, and co-registration and affine transformation of the functional volumes to corresponding T1w and subsequently to MNI space. For further details of the pipeline, please refer to the online documentation: https://fmriprep.org/.

To estimate connectivity between AnG and hippocampus, following previous studies studying AnG and episodic retrieval (Bonnici et al., 2016; Tibon et al., 2019), we defined the AnG region of interest (ROI) as a sphere of 6 mm radius (equivalent to 33 voxels) with its center at the stimulation site (x=-43, y=66, z=38, (Vilberg & Rugg, 2008)). The hippocampal ROI was obtained from a medial temporal lobe atlas (Ritchey et al., 2015). ROI-ROI resting-state functional connectivity analysis was performed using the CONN toolbox (Whitfield-Gabrieli & Nieto-Castanon, 2012). Preprocessed functional data were first linearly detrended and a commonly used bandpass filter (0.008–0.09 Hz) was applied to isolate low-frequency fluctuations characteristic of resting-state fMRI and attenuate signals outside of that range. White matter and CSF confound were removed using the aCompCor method. To ensure no voxels were included in mean estimates from outside ROIs, we performed all analyses using unsmoothed functional data.

## Results

### Vividness efficiency is causally dependent on angular gyrus

While it is often assumed that both vividness ratings and confidence ratings during retrieval mediate subjective experience of remembering, our primary aim was to test whether these two components of subjective mnemonic experience during retrieval are dissociable. We operationalized this idea by developing a paradigm, in which participants watched movies at encoding and performed a memory test immediately after receiving TMS inhibition to the AnG (Fig. 1, see Materials and Methods for details). In the memory test, participants mentally replayed relevant scenes with an image cue and rated the vividness of their memory. Following the vividness rating, participants were asked to make a temporal proximity judgment related to the image cues and rated the confidence of their memory judgment. Importantly, the temporal proximity judgments demand participants to mentally replay the cue related scenes for successful memory retrieval, thus we are confident that vividness and confidence ratings were to be made on the same memory traces. The novel and critical manipulation in our experiment is that subjective evaluation efficiency computed by vividness rating and confidence rating are differentiable under the recollection of the same segment of memory within a trial.

We first examined the effect of TMS to AnG on basic performance. As expected, TMS did not influence objective memory performance as measured by memory sensitivity d’ (mean_AnG_=0.79, *SD*_AnG_=0.23; mean_vertex_=0.90, *SD*_vertex_=0.35; t_19_=1.39, p=0.18, Cohen’s *d*=0.38; Fig. 2A) and reaction time (mean_AnG_=2.78s, *SD*_AnG_=0.43; mean_vertex_=2.84s, *SD*_vertex_=0.44; t_19_=0.68, p=0.51, Cohen’s *d*=0.14; Fig. 2B). Moreover, a repeated-measures ANOVA with subjective rating type (vividness/confidence) and TMS site (AnG/Vertex) for mean levels of subjective rating (confidence rating: mean_AnG_=2.85, *SD*_AnG_=0.32; mean_vertex_=2.84, *SD*_vertex_=0.42; vividness rating: mean_AnG_=2.79, *SD*_AnG_=0.31; mean_vertex_=2.76, *SD*_vertex_=0.43) did not reveal any significant main effects (rating type: F_(1,19)_=1.60, p=0.22, η^2^=0.08; TMS: F_(1,19)_=0.13, p=0.72, η^2^=0.01) nor an interaction (F_(1,19)_=0.21, p=0.66, η^2^=0.01; Fig. 2C). Of importance, we assessed whether inhibitory rTMS to left AnG modulated the efficiency of subjective ratings during memory retrieval (meta-efficiency: mean_AnG_=1.26, *SD*_AnG_=0.66; mean_vertex_=1.29, *SD*_vertex_=0.72; vivid-efficiency: mean_AnG_=-0.05, *SD*_AnG_=0.70; mean_vertex_=0.40, *SD*_vertex_=0.35) using two robust indices (vivid-d’/d’ and meta-d’/d’, see Materials and Methods). A repeated-measures ANOVA with factors of subjective efficiency type (vividness efficiency/metacognitive efficiency) and TMS site (AnG/vertex) revealed a significant main effect of efficiency type (F_(1,19)_=69.23, p<0.001, η^2^=0.78), as well as an interaction (F_(1,19)_=5.88, p=0.02, η^2^=0.24; Fig. 2D). Follow-up *t* tests revealed that participants showed significantly lower vividness efficiency following TMS to left AnG compared to vertex (t_19_=2.96, p_holm_=0.016, Cohen’s *d*=0.80), whereas no analogous decrement was found in metacognitive efficiency (t_19_=0.12, p_holm_=0.91, Cohen’s *d*=0.04). To better characterize the effect of AnG stimulation on vividness, we performed a sign test to verify the extent of changes between TMS to AnG and vertex. Reductions in vividness efficiency were consistent across participants due to TMS to AnG (16/20 reduced; sign test: p<0.001; Fig. 2E).

**Figure 2.**
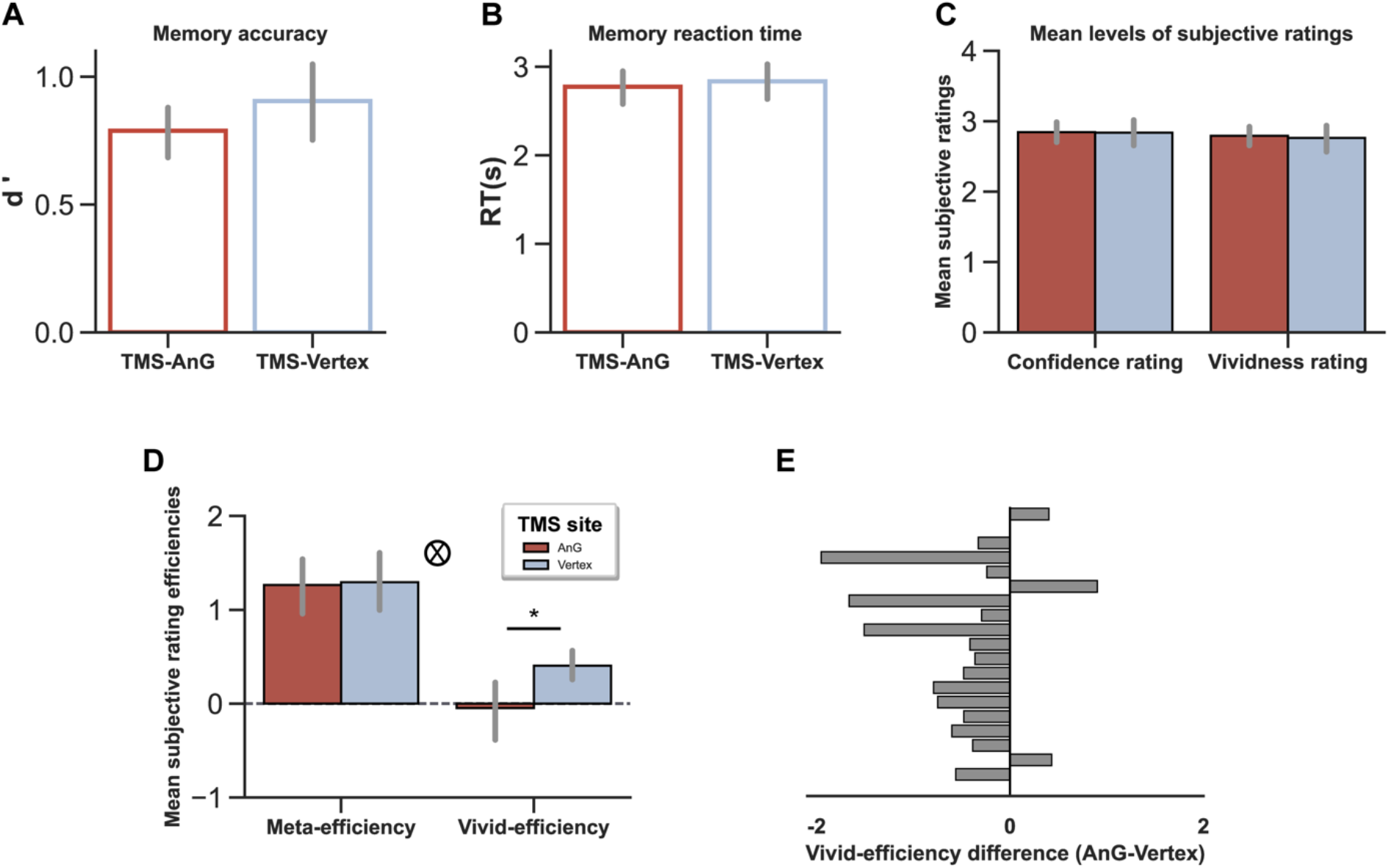
TMS effect on behavioral performance. **(A)** Accuracy (d’) and **(B)** Reaction times (RTs) in the temporal proximity task. **(C)** Mean levels of confidence ratings and vividness ratings. **(D)** Metacognitive efficiency and vividness efficiency. **(E)** Change in vividness efficiency between AnG and vertex stimulation for each participant. Error bars represent SEM. ^®^ indicates interaction of subjective reports efficiency by stimulation site in a repeated-measures ANOVA. *p < 0.05.

We further queried whether the vividness rating was reliably related to the temporal proximity memory performance using two additional analyses. First, we performed a permutation test to ensure the internal validity of the vividness efficiency index. Specifically, we randomly shuffled the vividness rating under TMS to vertex and re-calculated the vividness efficiency score for each participant (permutation n=1,000 per subject). Statistical significance was determined by comparing the true vividness efficiency to the null distribution of permutations for each participant. This analysis revealed that the vividness efficiency robustly quantifies the correspondence between vividness and objective memory in every participant (all p-values<0.005). Second, we assessed the efficiency of vividness on a trial-by-trial basis and tested the AnG TMS effect using a mixed-effects logistic regression model for objective memory performance against vividness ratings with the participant as a random effect for each stimulation site. Consistent with the observed TMS effect on vividness efficiency, we found that the vividness rating was a significant predictor of memory performance under TMS to vertex (β=0.213, p<0.001), but not under the AnG TMS condition (β=0.054, p=0.761). These two analyses show vividness efficiency, albeit its relatively low value, is a valid indicator for memory performance, both as a trial-wise and as a whole measure.

Subjective judgments (mainly metacognitive judgments) have been shown to exert a causal impact on the choice to collect more information (Desender et al., 2018; Metcalfe & Finn, 2008). We next asked whether the AnG TMS would impose any effect on the trade-off between memory accuracy and speed (RT). To do so, we computed an inverse efficiency score (mean correct RTs/% correct) to index the speed-accuracy trade-off. We did not observe a significant TMS effect on this speed-accuracy efficiency score (t_19_=0.30, p=0.76), suggesting that the observed TMS effect on vividness efficiency could not be explained away by any speed-accuracy tradeoff. In addition, to test whether the vividness ratings might be biased by the order of the target scene, we applied a 2 (TMS: AnG, vertex) × 2 (occurrence order: before, after) ANOVA to vividness efficiency and we found no significant interaction (F_(1,19)_=0.99, p=0.33, η^2^=0.05). This confirmed that the vividness ratings were not affected by the location of the target scene within the recalled segment. Moreover, to verify the lasting effects of TMS, we split the AnG TMS data into two halves based on their time within the experiment (first-vs. second-half). To test whether the observed TMS effect was modulated by time, we re-ran the vividness efficiency analysis for each half and submitted the vividness efficiency to a 2 (TMS: AnG, vertex) × 2 (Time: first-half, second-half) repeated-measures ANOVA. This revealed no main effect of Time (F_(1,19)_=0.002, p=0.96, η^2^<0.001) and no interaction involving Time (F_(1,19)_=1.22, p=0.28, η^2^=0.06), suggesting that the TMS effect did not differ in the first or second half of the experiment.

Together, these results suggest that the AnG is engaged in the monitoring of vividness and there might be a dissociation between vividness efficiency and confidence efficiency during episodic retrieval.

### AnG stimulation altered the mediating role of vividness in confidence in the accuracy of memory judgment

To examine how objective memory accuracy and the two subjective ratings of memory are interrelated in a single statistical framework, we conducted a mediation analysis using objective memory performance as the independent variable and vividness rating as the mediator variable under each TMS condition separately. We hypothesized that the link between objective memory response and confidence might be mediated by the vividness of memory. Under TMS to vertex, as expected, objective memory performance was significantly associated with both vividness ratings (ß=0.17, p<0.001) and confidence ratings (ß=0.56, p<0.001), indicating that both subjective ratings are meaningful in tracking the success of the same memory judgments. This is important because it allows us to test for the dissociation between vividness and confidence under the same TMS intervention. After adding vividness ratings as a simultaneous predictor, the relationship between objective memory performance and confidence ratings remained intact (ß=0.50, p<0.001). The trial-wise mediation analysis revealed that vividness ratings partially mediated the association between objective memory performance and confidence ratings (indirect effect = 0.06, p<0.001, 95% CI=0.04-0.08; Fig. 3A). Most importantly, by contrast, the vividness ratings did not mediate the relationship between objective memory and confidence ratings following AnG stimulation. The AnG stimulation altered the association between objective memory performance and vividness ratings (ß=0.05, p=0.149, Fig. 3B). Neither the relationship between confidence ratings and memory performance (ß=0.52, p<0.001) nor the relationship between confidence and vividness ratings (ß=0.33, p<0.001) was interrupted by AnG TMS. There was also a significant association between vividness ratings and confidence ratings (ß=0.34, p<0.001), which was not affected by AnG TMS. These findings indicate that, although both vividness ratings and confidence ratings were independently associated with objective memory performance under control site stimulation, AnG stimulation selectively impacted the association between vividness ratings and objective memory. These results provide further support to our main results (see Figure 2D and E) and revealed a mediation between memory performance and confidence through the subjective vividness of memory.

**Figure 3.**
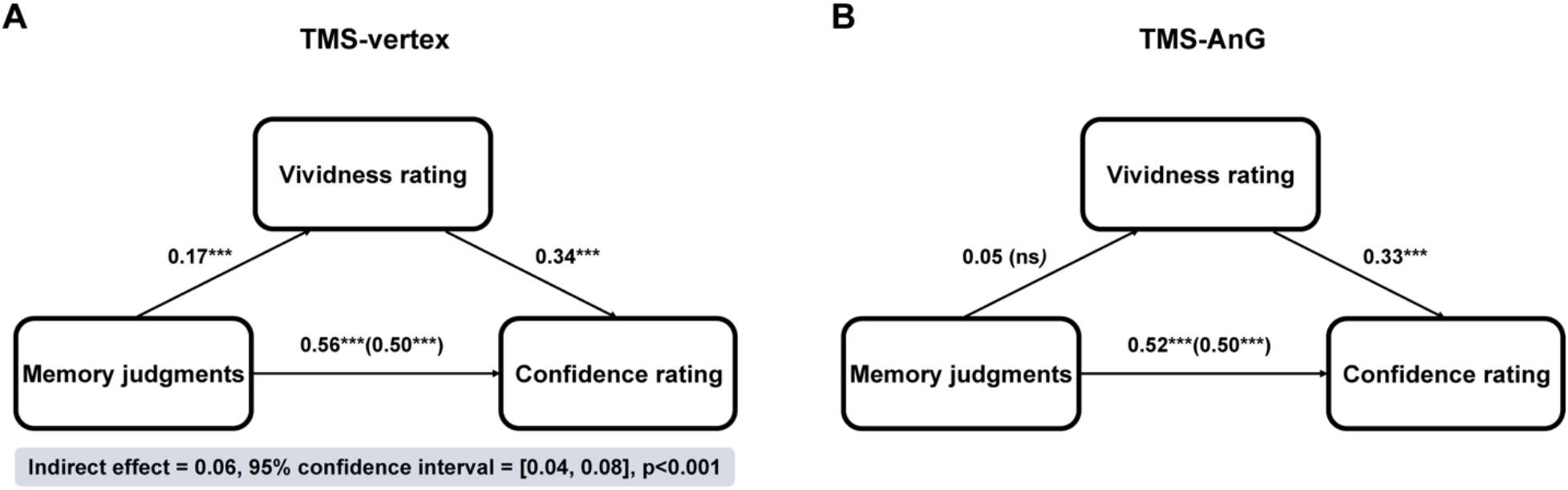
Mediation analysis between TMS conditions. **(A)** The mediation path diagram (vertex TMS condition) shows significant relationships between memory performance and vividness ratings; vividness ratings and confidence ratings; memory performance and confidence ratings; and a significant mediation effect of vividness on the relationship between memory performance and confidence ratings. **(B)** AnG TMS altered the association between objective memory performance and vividness ratings, while leaving the relationship between vividness ratings and confidence ratings; memory performance and confidence ratings unimpacted. ***p<0.001; ns = not statistically significant.

### AnG stimulation eradicated serial dependence effect in both subjective ratings RTs

We have thus far revealed differential TMS effects on the accuracy of two subjective ratings and their interrelationship with objective memory performance. We next sought to investigate whether the subjective evaluation mechanisms might share similarity in terms of how they incorporate past information into the current decision, or otherwise known as serial dependence effect (Fischer & Whitney, 2014; Rahnev et al., 2015, 2020). Given that RT is a defining element of the trade-off between speed and accuracy that characterizes decisions, the presence of serial dependence on RT can provide important insights into the nature of subjective awareness generation. To test for serial dependence in vividness RTs and confidence RTs separately, we performed a series of mixed regression analyses predicting subjective rating RTs with fixed effects for the recent trial history up to seven trials back and random intercepts for each participant. We also explicitly tested for any different involvement of AnG in generating subjective estimation during memory retrieval. We found that there was autocorrelation in vividness RTs up to lag-3 (all p-values < 0.05; Fig. 4A) under TMS to vertex. Following TMS to AnG, such serial dependence was not found any more. Furthermore, we also observed autocorrelation in confidence RTs up to lag-2 (all p-values < 0.05; Fig. 4B) under TMS control condition and such serial dependence effect was also reduced by AnG stimulation. These results replicated the existence of serial dependence in confidence RT and revealed serial dependence in vividness rating RTs, and both are modulated by AnG stimulation. The findings of such serial spill-over bias in both subjective estimations and their susceptibility to AnG stimulation might suggest their similarity in terms of subjective experience generations during memory retrieval.

**Figure 4.**
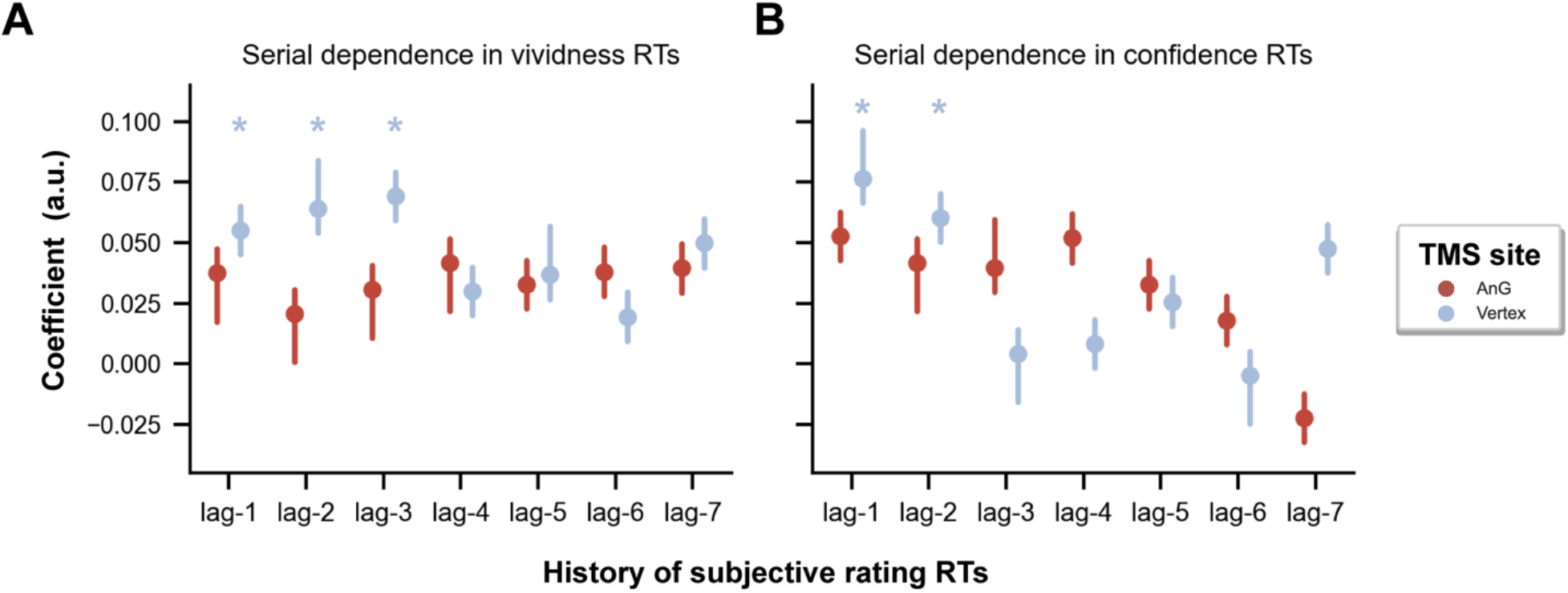
Serial dependence in subjective reports RTs. **(A)** Autocorrelation in vividness RTs was observed up to lag-3 under TMS to vertex (all p-values < 0.05; blue dots). No reliable autocorrelation was found in vividness RTs after TMS to AnG (red dots). **(B)** Autocorrelation was found in confidence RTs up to lag-2 under TMS to vertex (all p-values < 0.05). Such autocorrelation in confidence RT was also not found after TMS to AnG. *p<0.05.

### Resting-state functional connectivity between hippocampus and angular gyrus specifically relates to vividness efficiency

Having demonstrated that the AnG modulated the efficiency of vividness ratings, we then explored whether the intrinsic functional communication among brain regions was associated with subjective reports efficiencies. Specifically, we examined the relationship between intraindividual variability in subjective report efficiency and the resting-state functional connectivity between the AnG and hippocampus (Fig. 5A). This two regions has previously been shown to be related to memory metacognition (Baird et al., 2013). Interestingly, we observed a dissociation in this functional connection between efficiency of vividness and confidence. The functional connectivity of AnG-hippocampus was significantly correlated with vividness efficiency (r=-0.72, p<0.001; Fig. 5B), but not metacognitive efficiency (r=-0.27, p=0.243; comparison between correlations: z=1.876, p=0.03), suggesting that the vividness and confidence during memory retrieval may be mediated by distinct neural substrates. Moreover, TMS to AnG reduced the correlation between functional connectivity of AnG-hippocampus and vividness efficiency (r=-0.31, p=0.189; comparison between TMS sites: z=3.201, p=0.001). Consistent with our prediction, these results revealed that the self-monitoring of vividness and confidence are not only functionally but also neurally dissociable.

**Figure 5.**
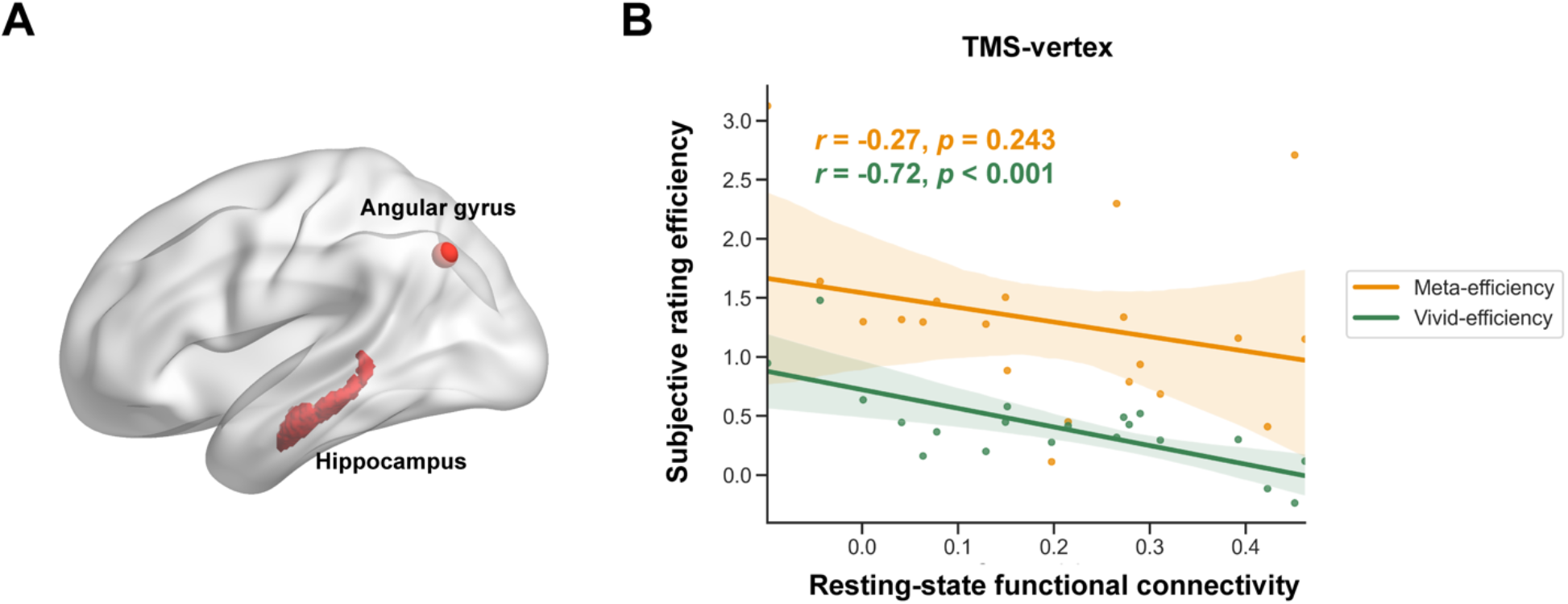
Resting-state functional connectivity analysis and anatomical double dissociation between the two subjective efficiencies. **(A)** ROIs (hippocampus and AnG). **(B)** Vividness efficiency, but not metacognitive efficiency, is significantly correlated with AnG-hippocampal functional connectivity.

In sum, consistent with our predictions, these findings establish a specific role of AnG, its mediating effects, and its functional connection with the hippocampus in subserving our perceived vividness of memory retrieval. The direct comparison with the metacognitive counterpart (indexed by confidence ratings) suggested functional and anatomical dissociation between the two subjective efficiencies in these mnemonic processes.

## Discussion

How do we obtain accurate assessment of our memory performance? Much of what we know about subjective aspects of memory comes from experimental work measuring the relationship between the level of confidence or vividness rating and neural activity during memory retrieval. Yet the ability to accurately monitor subjective mnemonic experience has remained poorly understood. Here, we asked the question of whether subjective confidence and vividness of memory reflect distinct introspective capacities. By administering non-invasive pre-retrieval stimulation to the left AnG, a candidate region supporting the subjective components of memory (Humphreys et al., 2021), we provide evidence for a causal involvement for AnG specifically in vividness efficiency. Critically, we show evidence that the ability of monitoring vividness of memory is indeed functionally and anatomically dissociable from confidence during episodic memory retrieval.

One of the novel aspects of this work is that we isolate the processes underlying vividness and confidence reports during episodic memory retrieval. We observed that temporary disruption of the AnG leads to difference in the efficiency of vividness ratings while leaving the efficiency of confidence ratings intact, suggesting that vividness and confidence of memory are two separable subjective experiences. These results are compatible with prior findings that the AnG is involved in the subjective experience of remembering (Kuhl & Chun, 2014; Yazar et al., 2014) but not in confidence-related metacognition. One possibility is that vividness of memory reflects something akin to the perception of past events, analogous to the ‘attention to memory’ (AtoM) account (Cabeza, 2008; Ciaramelli et al., 2008; Hutchinson et al., 2009). Retrieval from long-term memory demands selection between specific memories competing for recall (Badre et al., 2005). Previous theories have advanced the analogies between selection in the perceptual domain and selection during memory retrieval (Cabeza, 2008; Wagner et al., 2005). Accordingly, the AtoM account proposes that the parietal mechanisms (including AnG) support goal-directed attention toward the maintenance of mnemonic cues as well as facilitate the monitoring of episodic memory retrieval (Kwok & Macaluso, 2015; Hutchinson et al., 2009). In light of this view, the subjective sensed vividness during memory recall may thus represent a product of internal attentional processes rather than a subjective evaluation of memory quality, such as confidence. It is then plausible that the TMS to the AnG disrupts the shifting and allocation of attention to internal representations, resulting in less accurate perceived vividness of memory. A potential future direction following this work is to examine the degree of anatomical and functional convergence between the vividness rating and reflective attention.

Previous studies have linked activity in AnG with rated vividness (Bonnici et al., 2016; Kuhl & Chun, 2014) and reported that patients with lateral parietal lesions show diminished vividness or confidence of their memories (Berryhill et al., 2007; Hower et al., 2014; Simons et al., 2010). In the same vein, some other TMS studies have reported that AnG stimulation reduced confidence ratings of memory (Wynn et al., 2018; Yazar et al., 2014). Here, however, we did not observe any TMS effect on the overall reported vividness or confidence. One explanation for the discrepancy is that in our study, the participants encoded a naturalistic story per session and made memory judgments about the temporal proximity of two scenes, whereas in previous studies they used words and recognition task. As noted in Yazar et al. (2014), the observed TMS effect on mean confidence rating was specific to source recollection, while having cued recall confidence unimpaired, suggesting that differences in the types of task and stimuli might be responsible for producing differential AnG stimulation effects on mean subjective ratings across studies. Rather, instead of using the reported vividness, here we applied the concept of using performance and confidence correspondence (a quantitative measure of metacognition) to derive the degree of correspondence between rated vividness and objective memory accuracy. This approach enables us to estimate the TMS effect on the vividness efficiency independently from the level of vividness and objective memory performance. We asked participants to rate the vividness of the mental replay before any mnemonic decision, which allows for an uncontaminated assessment of the richness of mental experience prior to any memory judgement (Siedlecka et al., 2016). Our findings add to this limited literature by demonstrating a causal role for the AnG in vividness efficiency. One interpretation of these results is that the AnG may act as an accumulator in service of mnemonic decisions (Wagner et al., 2005). It has been previously proposed that memory retrieval is accomplished by a diffusion process during which evidence for a memory decision is accumulated (Ratcliff, 1978), and the parietal cortex, including AnG, is thought to play a role in the integration of sensory information (Gold & Shadlen, 2007; Shadlen & Newsome, 2001). This hypothesis is compatible with our data, accommodating the finding that TMS to AnG affected the correspondence between vividness and memory performance, but not the mean level of rated vividness and objective memory performance. Our findings clarify a role for AnG to accurately gauging the vividness of memory and support the notion that AnG participates in accumulating and integrating information in support of mnemonic processes.

In addition, intrinsic individual differences in functional connectivity between brain structures have informed our understanding of the varied ability to introspect about self-performance (Baird et al., 2013; Fleming et al., 2010; Ye et al., 2019). Here we found that resting-state functional connectivity of hippocampus and AnG is specifically associated with vividness efficiency, but not metacognitive efficiency, across individuals. This dissociation between functional connections between vividness and confidence efficiency is in line with our behavioral results that vividness and confidence may depend on dissociable neural substrates, suggestive of a differentiation account of subjective assessments of memory by functionally and anatomically dissociating the monitoring of vividness from confidence. A number of studies have showed AG is causally involved in episodic memory tasks (Bridge et al., 2017; Tambini et al., 2018; Wang et al., 2014). Regarding the influence of AnG TMS, we found that the relationship between vividness efficiency and AnG-hippocampal connectivity was eliminated under AnG TMS, consistent with the notion that the AnG TMS would distally modulate the function of hippocampus for memory processes (Wang et al., 2014). To put the results into a broader perspective, AnG is a key node within the default mode network, a set of brain regions that are consistently activated during rest, and deactivated during task (e.g. Buckner et al., 2008; Fox et al., 2005; Raichle et al., 2001). Interestingly, our finding revealed a negative relationship between vividness efficiency and AnG-hippocampal resting-state functional connectivity under vertex TMS. The negative correlation might be consistent with the proposal that suppression of the default mode network (including the AnG) is critical to success in some cognitive task performance (Anticevic et al., 2010). Future work combining TMS with fMRI could be used to examine to what extent TMS to AnG affect the interconnection between AnG and hippocampus during subjective memory processes.

Further, we observed a phenomenon of serial dependence in both subjective memory measures. These results extend previous demonstration of serial dependence in metacognitive judgments in perceptual tasks (Rahnev et al., 2020) to vividness and confidence judgments in an episodic memory task, suggesting that this phenomenon might be represented in a generic, task-independent format. We also showed that such effect was modulated by AnG stimulation, suggesting that the impact of AnG inhibition might go beyond subjective evaluation related to memory strength alone. Future studies should test whether this serial dependence phenomenon is domain-general and what factors might affect serial dependence in subjective evaluation judgments. In the literature on perceptual metacognition, theories of confidence generation posit that the central processing of evidence leading to a perceptual decision also establishes a level of confidence (Fetsch et al., 2014; Sanders et al., 2016). Some argue that confidence rating is corrupted by a meta-level noise (Shekhar & Rahnev, 2018; De Martino et al., 2012). In contrast, it remains less studied for the origins of confidence in the context of episodic memory decisions. Here, in an elucidation of the relationship between vividness, confidence, and objective memory performance, we found that vividness mediates the association between confidence and objective performance. This indicates that the sensed vividness of memory is instrumentally used for the computation of confidence. Consideration of the relative contribution of subjective feeling of vividness in generating confidence, especially for naturalistic paradigms involving continuous streams of multisensory information and mnemonic experiences, is thus paramount. Although the issue of deriving the best model for memory confidence is not our focus here, we hope that our findings provide some new insights into the confidence generation in episodic memory decision for future work. A critical avenue for future studies is to exploit what other information beyond subjective vividness is being used for confidence generation in episodic memory.

In closing, we demonstrate the contribution of AnG to vividness processing in terms of its mediating effect, its regional (by TMS), and cross-regional connectivity characteristics (by resting-state MRI). These findings suggest conscious mnemonic experiences could be elucidated by taking memory vividness, their relationship with confidence, and their anatomical profile into consideration.

## Acknowledgements

This research received support from National Natural Science Foundation of China (Grant No. 32071060), Science and Technology Commission of Shanghai Municipality (Grant No. 201409002800), Open Research Fund of the State Key Laboratory of Cognitive Neuroscience and Learning, and Jiangsu Qinglan Talent Program Award (S.C.K.).

